# Reward produces learning of a consciously inaccessible feature

**DOI:** 10.1101/2019.12.13.876243

**Authors:** Xue Dong, Mingxia Zhang, Bo Dong, Yi Jiang, Min Bao

**Author notes:** **Corresponding author**: Min Bao, **Email**.

## Abstract

Reward has significant impacts on behavior and perception. Most past work in associative reward learning has used distinct visual cues to associate with different reward values. Thus, it remains unknown to what extent the learned associations depend on the consciousness. Here we resolved this issue by using an inter-ocular suppression paradigm with the monetary rewarding and non-rewarding cues identical to each other except for the eye-of-origin. Thus, the reward coding system cannot rely on the consciousness to select the reward-associated cue. Surprisingly, the targets in the rewarded eye broke into awareness faster than those in the non-rewarded eye. We further revealed that producing this effect required both attention and inter-ocular suppression. These findings suggest that the human’s reward coding system can produce two different types of reward-based learning. One is independent of the consciousness yet fairly consuming attentional resource. The other one results from volitional selections guided by top-down attention.

## Introduction

It has long been recognized that reward and punishment strongly modulate behaviors and perception (Proshansky & Murphy, 1942). In addition, recent studies have brought evidence of reward processing abnormalities in patients with mental disorders (Whitton, Treadway, & Pizzagalli, 2015). Therefore, reward is becoming an increasingly important topic in both cognitive neuroscience and clinics.

The effects of associative reward learning in many studies (Anderson, 2017; Della Libera & Chelazzi, 2006; Marx & Einhauser, 2015; Proshansky & Murphy, 1942; Raymond & O’Brien, 2009; Serences, 2008; Wilbertz, van Slooten, & Sterzer, 2014) can be epitomized by a phenomenon known as the law of effect (Thorndike, 1911). The law denotes that a response repetitively followed by a reward will be more readily to recur when the participants encounter the same stimuli and context. In other words, actions or perceptions are biased in favor of reward-associated stimuli. In most cases, participants can consciously realize the rule of reward and thus intentionally direct attentional resources to the related stimuli. Therefore, reward appears to always work with selective attention. Indeed, numerous studies have suggested that attention and reward coding are closely linked processes for stimulus selection. For example, attentional inhibition of distractors in a prime display is robust for highly (but not poorly) rewarded prime display in a negative priming study (Della Libera & Chelazzi, 2006). Furthermore, facial stimuli associated with learned reward value can survive from attentional blink, suggesting privileged processing of reward-associated stimuli (Raymond & O’Brien, 2009). Interestingly, later studies report that learned reward value can bias attention independent of strategy and volition (Anderson, Laurent, & Yantis, 2011; Hickey, Chelazzi, & Theeuwes, 2010). In addition, reward and attention also similarly bias visual perception in binocular rivalry tasks (Marx & Einhauser, 2015; Wilbertz et al., 2014).

Nevertheless, it is already known that neuromodulatory signals for rewards (and punishments) are released diffusively throughout the entire brain (Dalley et al., 2001; Schultz, 2000; Vickery, Chun, & Lee, 2011). Intuitively, the effects of rewards could be free from the constraint of attention and consciousness. Thus, a lingering possibility to be tested is that reward learning can be established independent of the consciousness and selective attention. Unfortunately, to our knowledge, this hypothesis cannot be easily proved by the previous work, because in most of them perceptually distinguishable visual cues are used to associate with different reward values. Therefore, the structure of experiments *per se* may be responsible for their observations that rewards teach or rely on selective attention to choose objects of behavioral significance among a set of candidates. The present study introduces a novel paradigm in which the participants cannot consciously differentiate the monetary rewarding and non-rewarding visual cues. This was realized by rendering the two visual cues identical to each other except for their eye-of-origin information. With this design, participants should not be able to consciously tell the difference between the two monocular cues (Wolfe & Franzel, 1988; Zhang, Jiang, & He, 2012). Therefore, they could not realize which cue was associated with a reward. Selective attention is thus not believed to produce any eye-specific learning effects in such a paradigm. Any eye-specific learning effects established over the reward-based training should be contributed from the reward coding system, but independent of the consciousness and selective attention. We first showed that the eye-specific reward learning could be observed during inter-ocular suppression using a continuous flash suppression (CFS) method (Fang & He, 2005; Tsuchiya & Koch, 2005). With five more experiments, we further examined the role of top-down attention in the learning process and at what circumstances the eye-specific effects were present or absent.

## Results

Our initial experiment was based on a b-CFS paradigm (Jiang, Costello, & He, 2010). Reward was only associated with the detection of an invisible target in one of the two eyes (Figure 1A-B), which allowed us to examine whether reward learning can be established independent of selective attention. Participants completed two reward training sessions; whereas no reward was given in the pre- and post-test. Since the targets were in the same appearances whether it was presented to the rewarded eye or non-rewarded eye, selective attention was not likely to modulate the processing of associative reward learning in an eye-specific way. Any eye-specific learning effect in this training paradigm should be independent of consciousness. For each session, the breakthrough ratio was calculated by dividing the number of trials with correct responses by the total trial count for each eye-of-origin condition, respectively.

**Figure 1.**
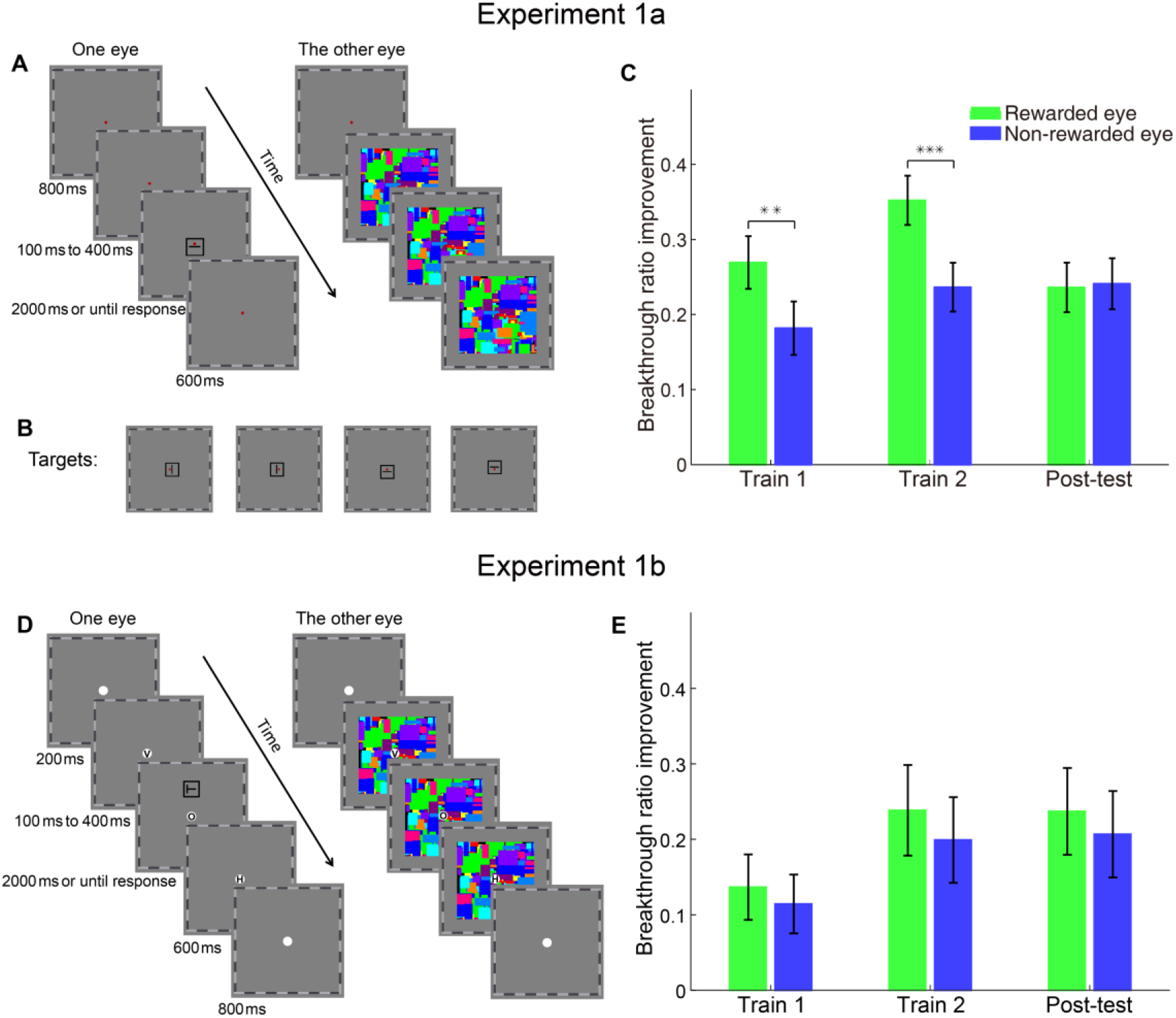
Stimuli and results of Experiment 1. (A), (B) Trial sequence and four kinds of targets in Experiment 1a. (C) The improvement of breakthrough ratio relative to pre-test in training and post-test sessions in Experiment 1a. (D) The stimuli and trial sequence in Experiment 1b. (E) The improvement of breakthrough ratio relative to pre-test in the training and post-test sessions in Experiment 1b. Error bars show ±1 SE. Asterisks indicate the significance level (with \*\**p* < 0.01, \*\*\**p* < 0.001).

Repeated measurements ANOVA and paired t-test were used to statistically analyze the results (see Table 1 for the detailed statistics of the ANOVA results). In the pre-test, there was no significant difference between the breakthrough ratios for the rewarded eye and the non-rewarded eye (*t*(35) = 0.193, *p* = 0.848, Cohen’s *d* = 0.028). Therefore, we subtracted the breakthrough ratios in the pre-test from those in the subsequent sessions to estimate the change of breakthrough ratios across the sessions. Consistent with the previous finding (Mastropasqua, Tse, & Turatto, 2015), the targets broke into awareness generally faster in the later sessions than in the pre-test. Surprisingly, there were more breakthrough trials in the rewarded eye than in the non-rewarded eye (training 1: *t*(35) = 2.815, *p* = 0.008, *d* = 0.463; training 2: *t*(35) = 3.649, *p* < 0.001, *d* = 0.658, see Figure 1C). However, this eye specific effect was absent in the post-test (*t*(35) = 0.438, *p* = 0.664, *d* = 0.059) where the participants no longer received monetary rewards.

**Table 1.** Statistics for the results of repeated measurements ANOVA.

|  |  | Before FA correction | After FA correction |
| --- | --- | --- | --- |
| Exp.1 | Session | $F(3, 105) = 49.768, p < 0.001, \eta^2 = 0.587$ | $F(3, 105) = 44.471, p < 0.001, \eta^2 = 0.560$ |
| | Eye-of-origin | $F(1, 35) = 3.785, p = 0.060, \eta^2 = 0.098$ | $F(1, 35) = 4.107, p = 0.050, \eta^2 = 0.105$ |
| | Session x Eye-of-origin | $F(3, 105) = 7.492, p < 0.001, \eta^2 = 0.176$ | $F(3, 105) = 7.348, p < 0.001, \eta^2 = 0.174$ |
| Exp.2a | Session | $F(3, 42) = 7.795, p < 0.001, \eta^2 = 0.358$ | $F(3, 42) = 6.849, p = 0.001, \eta^2 = 0.329$ |
| | Eye-of-origin | $F(1, 14) = 1.541, p = 0.235, \eta^2 = 0.099$ | $F(1, 14) = 1.828, p = 0.198, \eta^2 = 0.115$ |
| | Session x Eye-of-origin | $F(3, 42) = 1.293, p = 0.289, \eta^2 = 0.085$ | $F(3, 42) = 1.171, p = 0.332, \eta^2 = 0.077$ |
| Exp.3a | Session | $F(2.052, 34.878) = 28.364, p < 0.001, \eta^2 = 0.625$, Greenhouse-Geisser corrected | $F(2.037, 34.634) = 23.889, p < 0.001, \eta^2 = 0.584$, Greenhouse-Geisser corrected |
| | Eye-of-origin | $F(1, 17) = 0.293, p = 0.595, \eta^2 = 0.017$ | $F(1, 17) = 0.159, p = 0.695, \eta^2 = 0.009$ |
| | Target Orientation | $F(1, 17) = 9.774, p = 0.006, \eta^2 = 0.365$ | $F(1, 17) = 4.562, p = 0.048, \eta^2 = 0.212$ |
| | Target Orientation × Session | $F(3, 51) = 20.089, p < 0.001, \eta^2 = 0.542$ | $F(2.158, 36.689) = 10.466, p < 0.001, \eta^2 = 0.381$, Greenhouse-Geisser corrected |
| | Eye-of-origin × Target Orientation | $F(1, 17) = 3.209, p = 0.091, \eta^2 = 0.159$ | $F(1, 17) = 1.982, p = 0.177, \eta^2 = 0.104$ |
| | Eye-of-origin × Session | $F(3, 51) = 1.161, p = 0.334, \eta^2 = 0.064$ | $F(2.139, 36.371) = 0.941, p = 0.405, \eta^2 = 0.052$, Greenhouse-Geisser corrected |
| | Session × Eye-of-origin × Target Orientation | $F(2.049, 34.831) = 0.115, p = 0.896, \eta^2 = 0.007$, Greenhouse-Geisser corrected | $F(1.932, 32.847) = 0.173, p = 0.834, \eta^2 = 0.010$, Greenhouse-Geisser corrected |
| Exp.3b | Session | $F(4,60) = 16.782, p < 0.001, \eta^2 = 0.528$ | $F(4,60) = 15.438, p < 0.001, \eta^2 = 0.507$ |
| | Eye-of-origin | $F(1,15) = 1.916, p = 0.187, \eta^2 = 0.113$ | $F(1,15) = 1.676, p = 0.215, \eta^2 = 0.101$ |
| | Target Orientation | $F(1,15) = 0.073, p = 0.791, \eta^2 = 0.005$ | $F(1,15) = 0.076, p = 0.786, \eta^2 = 0.005$ |
| | Target Orientation × Session | $F(4,60) = 2.491, p = 0.053, \eta^2 = 0.142$ | $F(4,60) = 1.101, p = 0.365, \eta^2 = 0.068$ |
| | Eye-of-origin × Target Orientation | $F(1,15) = 1.258, p = 0.280, \eta^2 = 0.077$ | $F(1,15) = 0.954, p = 0.344, \eta^2 = 0.060$ |
| | Eye-of-origin × Session | $F(4,60) = 4.016, p = 0.030, \eta^2 = 0.211$,<br>Greenhouse-Geisser corrected | $F(4,60) = 4.021, p = 0.029, \eta^2 = 0.211$,<br>Greenhouse-Geisser corrected |
| | Session × Eye-of-origin × Target Orientation | $F(4,60) = 0.913, p = 0.462, \eta^2 = 0.057$ | $F(4,60) = 0.611, p = 0.656, \eta^2 = 0.039$ |

Here, FA denotes false alarm.

We further examined any potential influences of false alarms. In the present study, a false alarm represents that the participants made a wrong keypress in a trial. This could be due to a wrong judgment of the target position after the target broke into awareness or a cheating response by randomly guessing the location of the invisible target. In the training sessions, the participants might have the motivation to cheat in order to receive more monetary rewards. Considering the four-alternative forced-choice (4AFC) task, the probability of correctly guessed trials was 1/3 of that of wrongly guessed trials. Assuming that all the wrong responses were due to failure guesses, the maximum number of correctly guessed trials in theory could be estimated by dividing the number of trials with wrong responses by three. Therefore, to correct the effect of false alarm, the maximum number of correctly guessed trials was subtracted from the number of trials with correct responses before calculating the breakthrough ratios. After the correction, the breakthrough ratio also showed more increase in the rewarded eye than in the non-rewarded eye (training 1: *t*(35) = 2.782, *p* = 0.009, *d* = 0.465; training 2: *t*(35) = 3.562, *p* = 0.001, *d* = 0.653). The eye specific effect was absent in the post-test (*t*(35) = 0.329, *p* = 0.744, *d* = 0.044). In addition, by comparing the learning effect of the 1^st^ and 2^nd^ block of training 1, we found this eye-specific effect developed very fast in the training.

To test if top-down attention could affect the learning process, another group of participants were asked to perform a RSVP task simultaneously with the b-CFS task. The RSVP stimuli were presented on the center of the screen, and the b-CFS targets were presented 2° above or below the center of the screen. By using the fixation task and moving the b-CFS targets away from the central fixation position, the attention deployed on the b-CFS targets should be greatly reduced as compared to our initial experiment. Participants performed well in the RSVP task in all the sessions (hit rate: 87.23±7.85%, false alarm rate: 1.76±2.49%). The results of the b-CFS task, however, showed no significant difference in the breakthrough ratios between the two eyes in any of the four sessions (pre-test: *t*(13) = 0.637, *p* = 0.536, *d* = 0.178; training 1: *t*(13) = 0.764, *p* = 0.458, *d* = 0.145; training 2: *t*(13) = 1.141, *p* = 0.274, *d* = 0.180; post-test: *t*(13) = 1.615, *p* = 0.130, *d* = 0.141). Similar results were found after the false alarm correction.

### Monocular reward learning without inter-ocular suppression

Is inter-ocular suppression necessary for this eye-specific associative reward learning effect? To answer this question, we performed three experiments. We first tested the role of inter-ocular suppression using trials with and without CFS in Experiment 2a. Only in the trials without CFS (i.e. target-only trials) were participants rewarded for a correct response to the target presented to the rewarded eye. It was clear that these target-only trials were irrelevant to inter-ocular suppression. We found that in these trials, participants performed well in the pre-test for both eyes, with no significant difference in the performance (*t*(14) = 0.913, *p* = 0.377, *d* = 0.189). Training improved the performance for both eyes slightly by between 1.0% and 1.5% in the target-only trials with no difference across the eyes (all *p*s > 0.54). In the with-CFS trials, the break through ratio increased with training. However, no significant difference was observed between the breakthrough ratios for the rewarded eye and those for the non-rewarded eye in the pre-test (*t*(14) = 0.969, *p* = 0.349, *d* = 0.223), and the increase of breakthrough ratios did not show any difference between the two eyes in any of the training sessions (training 1: *t*(14) = 1.007, *p* = 0.331, *d* = 0.293; training 2: *t*(14) = 0.472, *p* = 0.645, *d* = 0.102; training 3: *t*(14) = 0.468, *p* = 0.647, *d* = 0.101, see Figure 2A). Similar results were found after the false alarm correction.

**Figure 2.**
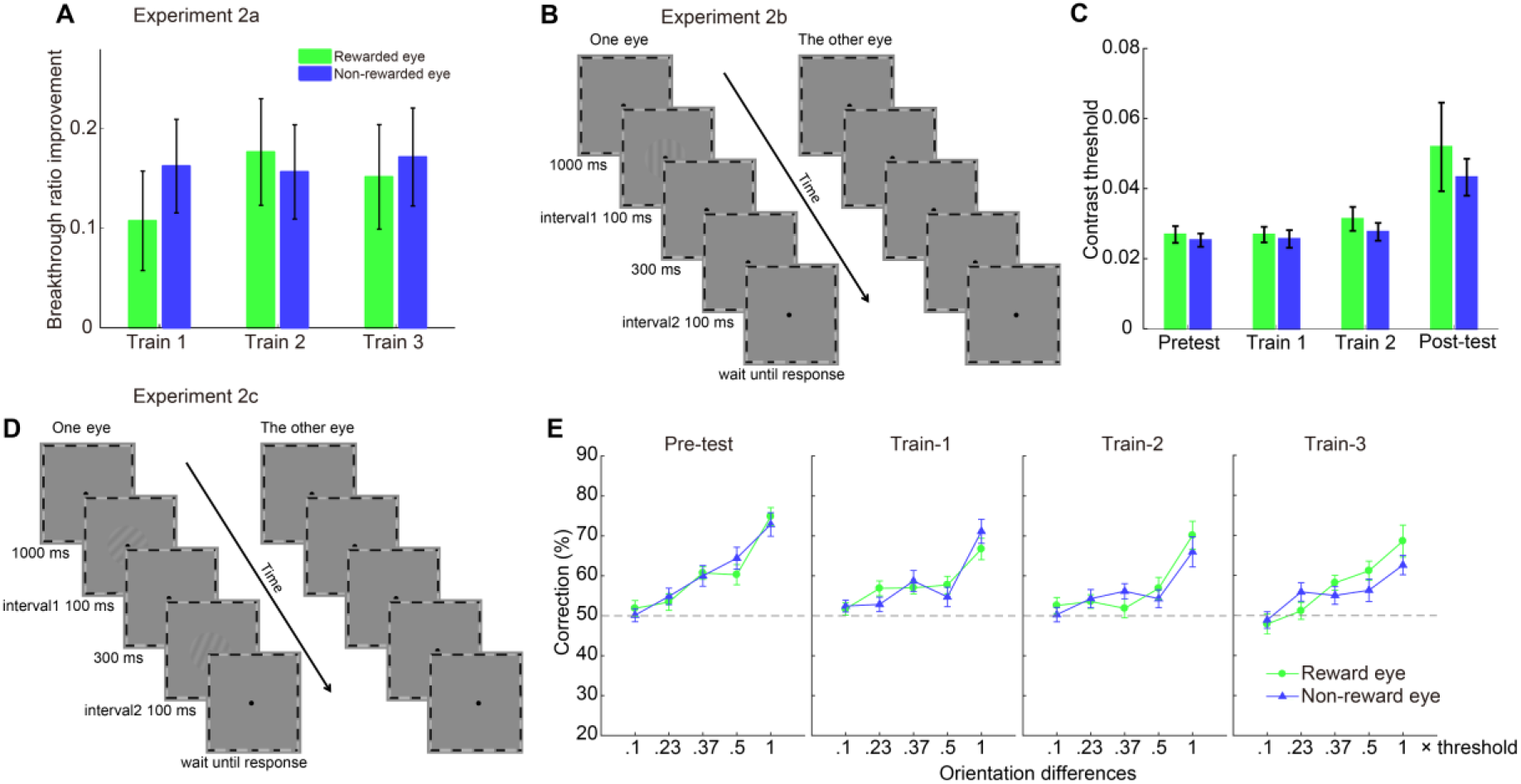
Stimuli and results of Experiment 2. (A) The improvement of breakthrough ratio relative to pre-test in three training sessions of Experiment 2a. (B) Trial sequence of the contrast detection task in Experiment 2b. (C) Contrast detection threshold of four testing sessions of Experiment 2b. (D) Trial sequence of the orientation discrimination task in Experiment 2c. (E) Orientation discrimination threshold of four testing sessions of Experiment 2c.

Since the performance for the target-only trials was almost perfect, a potential influence of ceiling effect could not be excluded. We then used a more difficult contrast detection task and an orientation discrimination task to further examine this issue. Subthreshold stimuli rather than inter-ocular suppression were used to render the stimuli hard to perceive in these two experiments.

In the contrast detection experiment (Experiment 2b, Figure 2B), no significant difference of the performance between the eyes was observed (Figure 2C, *F*(1, 13) = 1.986, *p* = 0.182, *η*^2^ = 0.133), though the contrast threshold showed a significant change across sessions (*F*(1.188, 15.446) = 7.977, *p* = 0.010, *η*^2^ = 0.380, Greenhouse-Geisser corrected). The main effect of Session was predominantly due to the increase of thresholds in the post-test than in other sessions, probably reflecting reduced motivation in the post-test that lacked incentives as compared to the training sessions.

Considering that the contrasts of stimuli in Experiment 2b were close to or below the detection threshold, visual signals to primary visual cortex might be faint. As a result, the eye-specific reward might not be able to enhance these signals. However, the eye-specific reward learning effect was still absent in Experiment 2c where high contrast gratings and orientation discrimination task were used (Figure 2D). For each offset level, the performances in the pre-test were not different between the two eyes (all *p*s > 0.80, FDR corrected). After subtracting the correction rates of the pre-test from those in training sessions, no difference was found between the rewarded and non-rewarded eyes on any levels in all the training sessions (Figure 2E, all *p*s > 0.19, FDR corrected).

### When selective attention was involved in the eye-specific rewarding

The results of the Experiments 1 and 2 indicated that inter-ocular suppression was necessary for eliciting the eye-specific reward learning effects when participants could not discriminate the rewarding vs. non-rewarding targets. In Experiment 1a, the eye-of-origin information was the only difference between the two kinds of targets, though could not be consciously realized. An interesting question is whether we can still observe the eye-specific learning effects when reward is also related to another feature that could be consciously distinguished. In Experiment 3, the rewarding target was defined by a conjunction of two features. Only the bar in one of the two orientations presented to the rewarded eye was the rewarding target. Although the participants were still not aware of the manipulation on the eye of origin information, they could quickly learn of the relationship between orientation and reward over training. We conducted Experiment 3 on two different groups of participants. Because of a mistake, the first group of participants did not complete the post-test after training (this was referred to as Experiment 3a hereafter). Therefore, we replicated the experiment in another group of participants but added the post-test (this was referred to as Experiment 3b hereafter). No significant differences among conditions were found in the pre-test of both groups (all *p*s > 0.18). Surprisingly, we found two different patterns of the learning effect in the two groups. Results from the first group of participants showed a significant interaction between Orientation and Session (see Table 1 for detailed statistics). Paired t-test indicated that the reward-based learning was orientation-specific (Figure 3B, RO-RE vs. NO-RE: training 1: *t*(17) = 1.336, *p* = 0.199, *d* = 0.176; training 2: *t*(17) = 2.687, *p* = 0.016, *d* = 0.621; training 3: *t*(17) = 4.205, *p* < 0.001, *d* = 0.861. RO-NE vs. NO-NE: training 1: *t*(17) = 1.559, *p* = 0.138, *d* = 0.164; training 2: *t*(17) = 5.092, *p* < 0.001, *d* = 0.457; training 3: *t*(17) = 6.864, *p* < 0.001, *d* = 0.785) but not eye-specific (*p*s > 0.20 for RO-RE vs. RO-NE and NO-RE vs. NO-NE). By contrast, results from the second group of participants showed a significant interaction between Eye-of-origin and Session. Further analysis revealed that the eye-specific learning effect became obvious in the last training session and even observable in the post-test where reward has been withdrawn (Figure 3C, RO-RE vs. RO-NE: training 1: *t*(15) = 1.307, *p* = 0.211, *d* = 0.314; training 2: *t*(15) = 0.961, *p* = 0.352, *d* = 0.302; training 3: *t*(15) = 2.359, *p* = 0.032, *d* = 0.827; post-test: *t*(15) = 2.587, *p* = 0.021, *d* = 0.691. NO-RE vs. NO-NE: training 1: *t*(15) = 0.307, *p* = 0.763, *d* = 0.077; training 2: *t*(15) = 0.934, *p* = 0.365, *d* = 0.223; training 3: *t*(15) = 1.343, *p* = 0.199, *d* = 0.421; post-test: *t*(15) = 2.013, *p* = 0.062, *d* = 0.599). Repeated measurements ANOVA disclosed a non-significant trend of interaction between Orientation and Session. An orientation-specific effect was only observed in the last training session in the rewarded eye, and was absent in the post-test (RO-RE vs. NO-RE: training 1: *t*(15) = 1.504, *p* = 0.153, *d* = 0.352; training 2: *t*(15) = 1.312, *p* = 0.209, *d* = 0.261; training 3: *t*(15) = 3.330, *p* = 0.005, *d* = 0.750; post-test: *t*(15) = 1.833, *p* = 0.087, *d* = 0.332; RO-NE vs. NO-NE: all *ps* > 0.16). Despite of two different learning patterns, the learning effects of both groups developed more slowly as compared to that in Experiment 1a. No significant learning effect was observed in the first training session. Similar results were found after false alarm correction.

**Figure 3.**
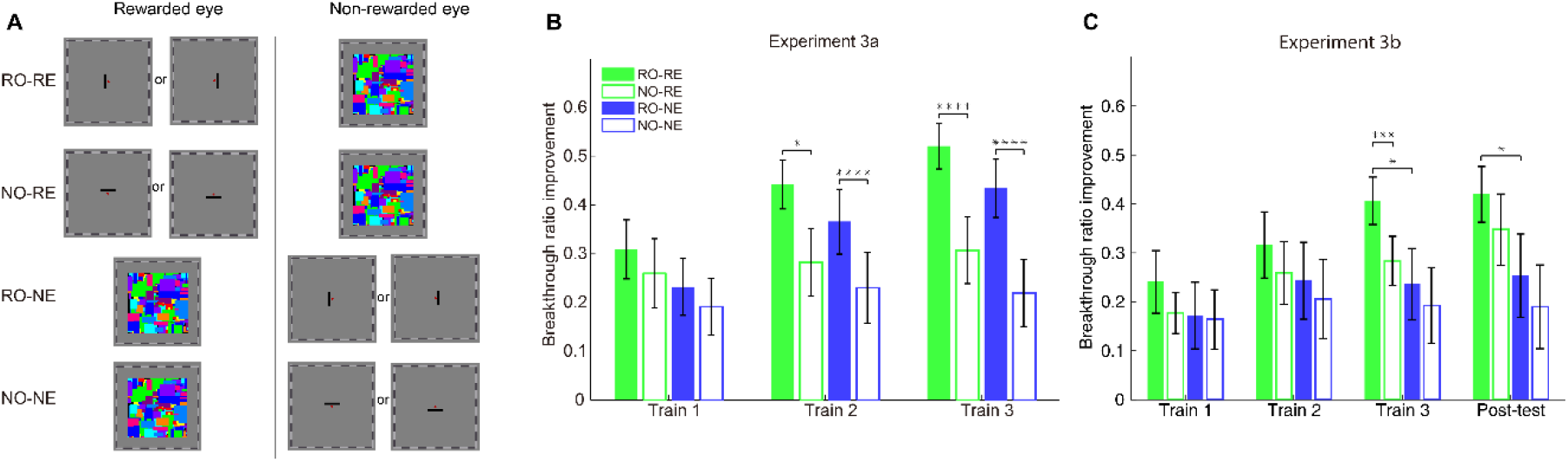
Stimuli and results of Experiment 3. (A) The examples of four kinds of trials in each session: rewarded orientation in the rewarded eye (RO-RE), non-rewarded orientation in the rewarded eye (NO-RE), rewarded orientation in the non-rewarded eye (RO-NE), non-rewarded orientation in the non-rewarded eye (NO-NE). The rewarded orientation and rewarded eye were counter-balanced across participants. (B)-(C) The improvement of breakthrough ratio relative to pre-test in training and post-test sessions of Experiment 3a and Experiment 3b. Asterisks indicate the significance level (with \**p* < 0.05, \*\*\**p* < 0.001, \*\*\*\**p* < 0.0001).

## Discussion

When a monocular pattern is presented to one eye, people have no explicit knowledge of the pattern’s eye of origin (Wolfe & Franzel, 1988; Zhang et al., 2012). We took advantage of this interesting nature of human vision to imbue the targets’ eye-of-origin information with monetary reward value. Because all the other features (e.g. shape, color, locations, etc.) of the targets were kept identical between the rewarding and non-rewarding conditions, the participants were not aware of the existence of two different types of targets. In Experiments 1 and 2, participants perceived a single kind of target accompanied with reward randomly in part of the trials in which they made a correct response. The importance of this design is that it dissociates consciousness and selective attention from reinforcement, thus allowing a close examination of the relationship between consciousness and reward coding.

The finding of an eye-specific learning effect in Experiment 1a supports our hypothesis that the reward coding system can rely merely on the eye-of-origin information of the reward-associated stimuli to produce a non-conscious reward-based learning effect. This result differs from Seitz and colleagues’ findings in their perceptual learning study by several aspects (Seitz, Kim, & Watanabe, 2009; Xue, Zhou, & Li, 2015). First, Seitz et al. used water as rewards for participants who were deprived of food and water (Seitz et al., 2009); whereas here the effects of monetary reward were investigated. Second, besides different eye-of-origin, the rewarding and non-rewarding control stimuli had different orientations in their experiment (Seitz et al., 2009); whereas in our Experiment 1a the only feature difference between the two stimuli was eye-of-origin. Third, the learning effect Seitz et al. found in their CFS experiment developed over 20 days of training, and could be observed in a separate sensitivity test where no reward was given (Seitz et al., 2009); whereas the effect in our Experiment 1a established quickly during the training and vanished immediately in a post-test that was conducted shortly after the training and with no reward. These distinct characteristics suggest that Seitz et al.’s findings, as they also proposed (Seitz et al., 2009), should be considered certain types of perceptual learning effects formed via a classical conditioning procedure. By contrast, the present findings remind us the brevity of non-conscious fear conditioning (Raio, Carmel, Carrasco, & Phelps, 2012). To our knowledge, the present study for the first time suggests that monetary reward, a positive and pleasant rather than a negative and threatening event (e.g. fear), can also induce non-conscious conditioning and the stimulus-reward pairing can be established only based on eye-of-origin—a consciously inaccessible feature.

In Raio et al.’s non-conscious fear conditioning study, the non-conscious fear learning was only significant during the early acquisition and declined quickly in the second half of trials, yet the conscious fear learning emerged slowly and became stronger during late acquisition (Raio et al., 2012). Similarly, we found that the reward learning could grow over time in Experiment 3 where the participants were aware that one orientation was more likely associated with rewards. Nevertheless, unlike Raio et al.’s finding, the non-conscious reward learning in our Experiment 1a did not show a rapid forgetting during the training. This might be due to different attentional status of participants in the training of the two studies. Raio et al. used a CFS paradigm in which participants did not have a task related to the suppressed stimuli (Raio et al., 2012); whereas we used a b-CFS paradigm so that participants had to pay attention and report any potential breakthroughs of the suppressed stimuli. Not only that, attention even seems to play an important role in generating the eye-specific reward-based learning effect, because the effect was absent when attention was distracted from the rewarding task. We speculate that top-down attention may in some way help discern the eye-of-origin information associated with rewards or even likely participates in reinforcing this association. However, the contribution of top-down attention here should be clearly different from the more common roles of selective attention in modulating the actions or perceptions when participants have an explicit knowledge of the rule of reward delivery in many previous studies (Della Libera & Chelazzi, 2006; Marx & Einhauser, 2015; Raymond & O’Brien, 2009; Serences, 2008; Wilbertz et al., 2014).

In the past decade, reward has frequently been reported to induce changes in early sensory regions (Arsenault, Nelissen, Jarraya, & Vanduffel, 2013; Chubykin, Roach, Bear, & Shuler, 2013; Liu, Coleman, Davoudi, Zhang, & Hussain Shuler, 2015; Persichetti, Aguirre, & Thompson-Schill, 2015; Pleger, Blankenburg, Ruff, Driver, & Dolan, 2008; Serences, 2008; Shuler & Bear, 2006; Zold & Hussain Shuler, 2015) but see (Rossi et al., 2017). The current work provides a strong case arguing that reward can induce plasticity in human V1 which is largely independent of consciousness. Such relatively lower level plasticity should be driven by the co-work of top-down eye-based attention (Zhang et al., 2012) and diffusively distributed neuromodulatory signals released by the reward coding system (Dalley et al., 2001; Schultz, 2000; Vickery et al., 2011). During inter-ocular suppression, the invisible rewarding target activates monocular neurons for the rewarded eye, while the activities of binocular neurons are mainly dominated by the signals for the CFS stimuli. Obviously, only the firings of monocular neurons for the rewarded eye are highly predictive of later rewards, rather than those of binocular neurons (and those of monocular neurons for the non-rewarded eye). During the training, the reward coding system may soon detect the reliable association between rewards and responses of monocular neurons for the rewarded eye. Eye-based attention may then increase the gains particularly for those neurons. Previous work has revealed that attention and consciousness can be independent to each other in some cases (Brascamp, Van Boxtel, Knapen, & Blake, 2010; Van Boxtel, Tsuchiya, & Koch, 2010). Therefore, it is not strange that in the present study attention can work with the reward coding system independent of consciousness. As a result, the breakthrough was facilitated more for the rewarded eye than for the non-rewarded eye in Experiment 1a. Once the attentional resource was consumed by another demanding task, the co-work of attention and reward coding system failed, thus no eye-specific learning effect was observed in Experiment 1b.

Notably, the above explanation receives further support from the results of our Experiment 2 where we examined whether inter-ocular suppression of the target was necessary for observing the eye-specific learning effects. The results of Experiment 2a showed no eye-specific learning effects. To avoid any unwanted influence of the ceiling effect, we used more difficult tasks in Experiment 2b and Experiment 2c. Again, no eye-specific learning effects were observed. Therefore, inter-ocular suppression seems to be a necessity for the finding in Experiment 1a. Without inter-ocular suppression during the training, the targets were represented by the activities of both monocular and binocular neurons. However, the relationship between rewards and neuronal responses was different in these two neuronal populations. In case of breakthrough, the firings of monocular neurons were either 100% (for the rewarded eye) or 0% (for the non-rewarded eye) predictive of subsequent rewards, yet the firings of binocular neurons were always 50% predictive of rewards. The absence of the eye-specific learning effects thus indicated that in Experiment 2 the reward coding system weighted heavily on the activities of binocular neurons and ignored the eye-of-origin information. This is possible given that binocular neurons greatly outnumber monocular neurons in the visual cortex.

Is inter-ocular suppression sufficient to observe the eye-specific learning effects? As indicated by our Experiment 3, this is not the case. As shown in Figure 3, one group of participants showed only the orientation-specific learning, while the learning of the other group of participants relied on both the orientation and the eye-of-origin information. In Experiment 3, a rewarding target was defined as a feature conjunction, so that both the orientation and eye-of-origin had a 50% probability to be a feature of the rewarding target, though the actual probability for the rewarding target was 25%. However, over training the participants might only realize that one of the two orientations was reward-associated, while still unaware of the relationship between rewards and eye-of-origin. As shown in Figure 3B, the first group of participants only showed the orientation-specific learning, although rewards were only preceded by half of the targets that were in the reward-associated orientation (i.e. RO-RE rather than RO-NE), they might treat both the RO-RE and RO-NE targets as a single type of targets that sometimes (50% probability) brought rewards at the time of breaking into awareness. As a result, targets in the reward-associated orientation, which were probably expected by the participants, were more readily to break into awareness than those in the perpendicular orientation. The increase of breakthrough ratio was thus similar for the two conditions. In this case, the reward coding system seemed to no longer work in an unsupervised mode based solely on the eye-of-origin information as in Experiment 1a. Instead, it worked in a supervised mode, and selectively strengthened the representations in the expected orientation. This behavior of the reward coding system is very similar to its role in teaching attention to make selections which is advocated by many other researchers. Importantly, this supervised-mode mechanism seems to override the unsupervised-mode mechanism, leading to an absence of the eye specificity. By contrast, the learning in the second group showed a more complex pattern. These participants learned even more slowly than the first group, and in the last training session they showed both an orientation-specific and eye-specific learning effect. We presume that the slower development of the orientation-specific learning may indicate a lower efficiency of the supervised-mode mechanism in these participants. Accordingly, the unsupervised-mode mechanism might have the chance to play a role, eventually resulting a significant eye-specific learning effect.

As compared to Experiment 1a, the more complicated reward rule in Experiment 3 might hamper a fast reward-based learning. Consistent with this conjecture, both groups of participants in Experiment 3 learned more slowly than the subjects in Experiment 1a. Alternatively, the sluggish learning in Experiment 3 might result from the additional use of a consciously accessible feature (i.e. orientation), whereas in Experiment 1a only the consciously inaccessible feature was used to associate with rewards. This also agrees with Raio and colleagues’ finding that participants learned more slowly when they were aware of the reward-associated stimuli than when not (Raio et al., 2012). Besides different temporal pattern of the learning between Experiments 1a and 3, we also noticed that the learning effect was immediately extinct in the post-test of Experiment 1a, yet still detectable in Experiment 3. Although illuminating this interesting diversity still awaits more systematic future work, we provide a bold surmise that the storage of a learned reward association may be strengthened by the consciousness, thus can be easier or only works when the participants consciously realize the reward-associated feature.

We therefore propose that the reward coding system can produce two different types of reward-based learning. One of them is independent of the consciousness yet fairly consuming attentional resource, likely occurring as early as in V1. We call it the unsupervised learning effect. The other type of learning rests on the close interactions between reward and selective attention, which we call the supervised learning effect. The supervised learning effect can override the unsupervised learning effect when voluntary attention participates in selecting stimuli of behavioral significance. In our Experiment 3a, this is probably because top-down attention selectively enhances the representations in the expected orientation regardless of their eye-of-origin information. With respect to the temporal nature, the eye-specific unsupervised reward learning effect is likely to develop very fast in the training (e.g. reaching the asymptote within the first block of the first training session in Experiment 1a), and highly depends on the context of reward delivery. As soon as the context of reward delivery was absent (e.g. in the post-test), the effect was also absent. Once the supervised reward learning joined, the temporal pattern of the learning changed, hinting distinct timescales between the two types of reward-based learning.

In summary, the present study reports a novel type of reward learning that is established specific to one eye. Physically identical targets were presented either to the rewarded eye or to the non-rewarded eye, yet rewards were delivered only after the targets in the rewarded eye broke into awareness. Over training, the breakthrough for the rewarded eye was facilitated more than that for the non-rewarded eye. This phenomenon was only reliably observed when the targets were rendered invisible using inter-ocular suppression and the only difference in the visual input signals between the rewarding and non-rewarding conditions was the eye-of-origin information. Our findings suggest that human’s reward coding system can produce an unsupervised reward learning effect independent of the consciousness.

## Materials and Methods

### Participants

Thirty-six participants (17 males and 19 females) who were screened from 105 volunteers participated in Experiment 1a. Fourteen participants (7 males and 7 females) who were screened from 45 volunteers finished Experiment 1b. Fifteen participants (5 males and 10 females) who were screened from 57 volunteers finished Experiment 2a. Two groups of fourteen participants finished Experiment 2b (6 males and 8 females) and Experiment 2c (4 males and 10 females). Eighty-three volunteers were recruited for Experiment 3 and 34 of them passed the screen test. Eighteen (12 males and 6 females) participated in Experiment 3a, 16 (5 males and 11 females) participated in Experiment 3b. The participants ranged in age from 18 to 28 y, all had normal or corrected-to-normal vision and were naïve to the experimental hypotheses. Experimental procedures were approved by the Institutional Review Board of the Institute of Psychology, Chinese Academy of Sciences, and informed consent was obtained from all participants.

### Apparatus

Stimuli were presented on a 21-in Dell CRT monitor with a resolution of 1024 × 768 pixels at a refresh rate of 85 Hz, and programmed in Matlab and Psychtoolbox 3 (Brainard, 1997). The display was calibrated with a Photo Research PR-655 spectrophotometer. To calibrate the display, we measured the luminance gamma curves and inverted them with a look-up table. The mean luminance of the screen was 50.9 cd/m^2^. A chin-rest was used to help minimize head movement.

### Stimuli and Procedures

#### Experiment 1

##### Experiment 1a

Stimuli were dichoptically presented on a mid-grey background. The target was a dark gray square frame (1.2° × 1.2°, linewidth: 0.11°) with a horizontal or vertical bar (length: 0.8°, linewidth: 0.11°) in the center. The target was displayed foveally in one eye, centered 0.25° away from the central fixation point (0.2°). The central bar could be vertically oriented to the left or right of the fixation point, or horizontally oriented above or below the fixation point (see Figure 1A). The CFS stimuli (8° × 8°, flashing at 10 Hz) were displayed foveally in the other eye, which consisted of 60 Mondrian patterned images created by drawing rectangles of random colors and sizes. A black-and-white square frame (11° × 11°, linewidth: 0.11°) and the central fixation point were always presented binocularly to help fusion.

Each trial started with a presentation of the central fixation point for 800 ms. Afterwards, the CFS stimuli appeared in one eye and kept flashing until the end of the trial. The target appeared in the other eye after a random interval varied between 100 and 400 ms. The contrast of the target ramped up to its highest level within the initial 1500 ms, and then remained at the highest contrast level for 500 ms. Participants were required to report the position of the central bar of the target relative to the fixation once the target broke into awareness. They were told to respond as quickly as possible on the premise of accuracy. The trial terminated once a response was made, otherwise the target would be displayed for 2000 ms in total followed by a 600-ms blank interval while the CFS stimuli were still presented in the other eye. To prevent any afterimage of the target, the target kept drifting back and forth for 0.08° at 1 Hz along the diagonal.

Each volunteer first participated in a screen test to find individuals with relatively balanced eyes and the stimuli contrast for non-extreme breakthrough ratio for each eye. Specifically, for targets of a certain contrast, the difference of break ratio across the eyes should not exceed 20%, while the break ratio for each eye had to be between 20% and 60%. We used this criterion (around 40%) in case the estimation of training induced increment of breakthrough ratio was affected by a floor effect or a ceiling effect from repeatedly performing a b-CFS task. The criterion was decided in a preliminary stage of the study. Eventually, the break ratios of 29/36 of the participants were within this range. For the other 7 participants, we used a less stringent criterion (18% - 65%) to have a larger sample. The contrasts for the target and CFS determined by the screen test were then used in the subsequent formal experimental sessions. The optimal contrasts used for each participant were listed in the Table S1, as well as the breakthrough ratios for each eye.

After the pre-test, the participants completed two training sessions, and a post-test. Each session consisted of two blocks of 160 trials. In the pre- and post-tests, there were no monetary rewards. The target was presented to the left eye in half of the trials, and to the right eye in the rest of the trials. The two conditions of trials were randomly interleaved. In the training sessions, for each participant one eye was assigned to be the rewarded eye. Participants were not aware of this setting. The selection of the rewarded eye was counter-balanced across the participants. A trial was called a rewarding trial if the target was presented to the rewarded eye. Immediately after a correct response for a rewarding trial, a 500-Hz tone would beep for 50 ms, which notified the participants of winning 0.2 yuan. After each block, a message on the screen showed the participants the total amount of gain.

##### Experiment 1b

The stimuli of b-CFS task were similar to those in Experiment 1a, except that the target was a capital letter “T” in a squared frame. The target was presented at 2° eccentricity above or below the center of the screen. The letter has four orientations (upright, upside down, right tilt, left tilt), participants were asked to report the letter orientation by pressing the corresponding arrow key. Simultaneously with the b-CFS task, participants were required to complete a central RSVP task. A series of capital letters were binocularly presented in a central white circle (0.6° in diameter) during the presentation of CFS stimuli. Each letter subtended for 0.5° and was presented for 250 ms. The task was to find “O” in the letter series. A trial lasted for 3700-4000 ms or until the press of an arrow key was detected. Only one letter “O” was presented in each trial, and it was not presented at the beginning or the last 200 ms. In each block, 20 trials were planned to be catch trials without RSVP target. However, since a trial may end before the presentation of “O”, there would be more catch trials. We calculated the hit rate and false alarm rate to measure the performance of RSVP task. A hit was the response that was made after the presentation of “O” and before the end of the trial. A false alarm was the response that was made before or without the presentation of “O” in a trial.

The screen test and procedure were same as those in Experiment 1a. In the training sessions, participants were told that the reward they could receive was firstly rely on the b-CFS task, while the hit rate of RSVP task would serve as a discount ratio to the overall reward.

#### Experiment 2

##### Experiment 2a

The stimuli and task were similar to Experiment 1a. However, there were two different types of trials in the training sessions, with-CFS trial and target-only trial. The with-CFS trial was identical to a typical trial in Experiment 1a. In a target-only trial, no CFS stimuli were presented, and a target ramped up from 0 to −0.8 contrast (Weber contrast, C = (L_s_ − L_b_) / L_b_, where L_s_ and L_b_ denoted the luminance of the stimulus and background) within 2 s in one of the two eyes.

The screen test and pre-test were the same as those in Experiment 1a. During the training sessions, one eye was assigned to be the rewarded eye. However, rewards just occurred in the *target-only* trials where the target appeared in the rewarded eye. A correct response in a rewarding trial would bring a 500-Hz beep and give rise to a reward of 0.31 yuan. In this experiment, all the participants could finish the task with nearly perfect performance. Therefore, in order to make the total amount of rewards slightly different across the participants, a random amount (ranging from −5.00 to 5.00 yuan) was added to the final gain for each block before it was shown on the screen. Each participant completed a pre-test and three training sessions.

##### Experiment 2b

The stimuli were sinusoidal gratings (3° in diameter, 1.5 cpd), presented monocularly on the center of a mid-grey background. The orientation of gratings was fixed for each participant (either vertical or horizontal), but was counter-balanced across participants.

Participants performed a two-interval forced-choice (2IFC) task. They were required to detect in which interval the grating was presented. Three practice sessions and four formal test sessions were completed. The contrasts of test gratings were manipulated by a 2-down-1-up staircase in the practice sessions. Sixty contrast levels were predetermined for the staircase, ranging logarithmically from 0.4% (Michelson contrast) to 4% (though 10% for the first practice session for an easier task). The first practice session included only one block, which was used for the participants to get familiar with the task. A block contained two interleaved staircases. Each staircase consisted of 60 trials and started with the highest contrast level. The test contrast decreased after two successive correct responses and increased after every wrong response. The step size for the staircase was initially three contrast levels and was reduced to one contrast level after three reversals. The procedure was similar for the other two practice sessions except that each session contained two blocks. Contrast threshold for each staircase was calculated by averaging the contrast levels of the last six reversals. The mean threshold of the latter two practice sessions was used to determine the seven contrast levels in the formal experiments, which were designed with the constant stimuli method. Similar to the formal experiments, feedback beeps (1200 Hz) were given after the response. However, they were delivered randomly in half of the trials with correct responses and participants were told to ignore the beep.

After the practice, participants finished four formal test sessions, a pre-test session, two training sessions and a post-test session. The task was same as that used in the practice experiment. Seven test contrast levels ranged logarithmically from one fourth to three times of the threshold estimated in the practice sessions were used. For each eye, every contrast level was tested 50 times, resulting in 700 trials per session. The test eye and contrast were randomly selected in each trial. A session was divided into 4 blocks, allowing the participants to take a break after every block. The contrast threshold of each eye was estimated by fitting the accuracies at all contrast levels with a Weibull function (82% correct performance).

Unbeknown to the participants, one eye was selected to be the rewarded eye before the formal experiments. The selection of rewarded eye was counter-balanced across participants. Every correct response for a rewarding trial was accompanied with an auditory feedback (1200 Hz). However, only in the training sessions, participants were informed that the high frequency beep meant an extra money reward of 0.77 yuan. After each training block, a message on the screen showed the participants how much they had earned.

##### Experiment 2c

The stimuli were gratings with the contrast of 80%, and the orientations were about 45°. In each trial, the stimuli were presented monocularly. Participants performed an orientation discrimination task by judging whether the second grating tilted clockwise or counter-clockwise to the first one.

After a few practice sessions, participants performed a pilot session where their orientation discrimination thresholds were measured. The orientation difference between the reference and test gratings was adjusted according to a staircase procedure. Each practice session included two interleaved staircases, one for each eye. The pilot session included four interleaved staircases, two for each eye. Every staircase contained 50 trials. Fifty levels of the orientation offset were predetermined for the staircase, ranging logarithmically from 0.1° to 10°. The mean orientation offsets from the last six reversals of each staircase were calculated as the orientation discrimination threshold for each eye respectively (71% correction threshold).

In the formal experiments, participants finished a pre-test session and 3 training sessions. Each session consisted of 5 blocks (100 trials per block). For each eye, four subthreshold offset levels (0.10, 0.23, 0.37, 0.5 × individual threshold) and a threshold offset level were determined. The threshold level was set to ensure participants could discrimination the orientation difference in some of the trials so that they would not give up on the task. In each block, 10 trials were tested for each offset level.

There was no reward in the pre-test. In the training sessions, one eye was selected as the rewarded eye. For trials in which stimuli with the maximum orientation offset (threshold level) were presented to the rewarded eye, a beep (1300 Hz) sounded immediately after the participants made a correct response. For stimuli with subthreshold orientation offset and were presented to the rewarded eye, the beep was given regardless of whether the responses were correct or not. Participants were not informed under what circumstance the reward would be given, but were instructed that each beep meant that they had earned a certain amount of reward (0.09 yuan/trial). The total gain was presented on the screen after each block. It should be noted that, since participants were rewarded according to the correction their responses for only 10 trials in each block, the amount of reward could be nearly equal across the blocks. To increase the variance of reward over blocks, a random value was either added to or subtracted from the actual money participants had earned.

#### Experiment 3

The stimuli were similar to those in Experiment 1a, except that the target was either a vertical or horizontal bar (length: 1.4°, linewidth: 0.2°) without the square frame (see Figure 3A). This could make a vertical bar more distinguishable from a horizontal bar.

Like Experiment 1a, participants were required to detect a target in one eye suppressed by the CFS stimuli in the other eye. Because there were two different targets (vertical or horizontal) and the target could be presented to one of the two eyes, there were four conditions in this experiment (vertical target in the left eye, vertical target in the right eye, horizontal target in the left eye, and horizontal target in the right eye). Only one of the four conditions (counter-balanced across the participants) was assigned to be the rewarding condition, while the other three conditions were non-rewarding conditions. A correct response in a rewarding trial would produce a reward of 0.5 yuan accompanied by an audio feedback. The gross of rewards was listed in a message on the screen after the end of each block. The four types of trials were randomly interleaved within a session.

Since the eye specific learning effect in Experiment 1a was no longer observed once the reward was withdrew in the post-test, we first asked a group of participants to complete the experiment with a screen test, a pre-test, and three training sessions (Experiment 3a). Another group of participants completed the experiment with an extra post-test without reward after training (Experiment 3b) to test the persistence of the learning effect.

## Acknowledgments

This research was supported by the National Natural Science Foundation of China (31871104, 31571112, 31500888, and 31830037) and the Key Research Program of Chinese Academy of Sciences (XDB02010003 and QYZDB-SSW-SMC030).

## Author Contributions

M.B. conceived the study; X.D., M.Z., Y.J. and M.B. designed the experiments; X.D., M.Z. and B.D. performed the experiments; X.D., M.Z. and M.B. analyzed the data; X.D. and M.B. wrote the paper.

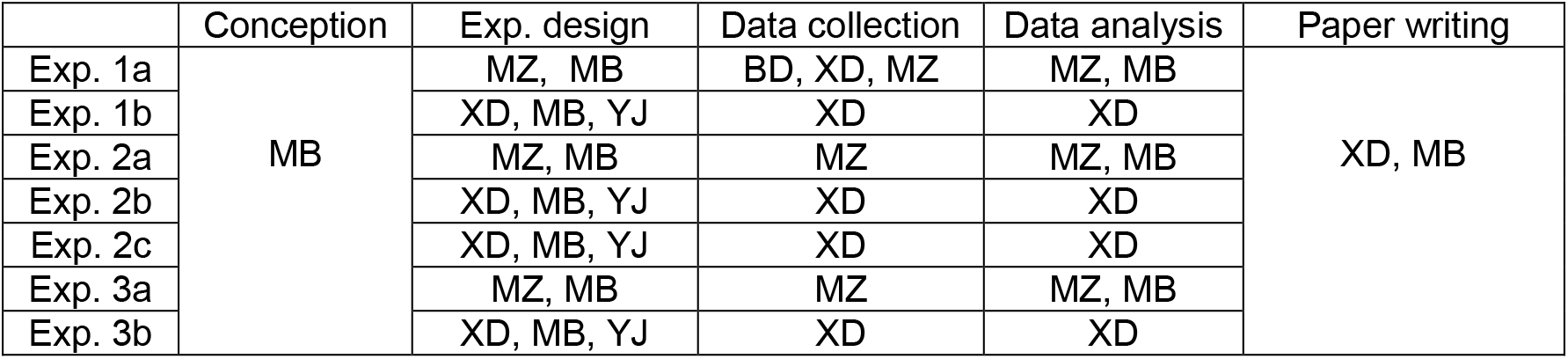

