## Supplemental Tables 1-5 for "Reward produces learning of a consciously inaccessible feature"

Supplemental Tables 1-5: Finally determined target contrast and the breakthrough ratio at the determined contrast for subjects who completed the whole procedure in the pretest of Experiment 1a, Experiment 1b, Experiment 2a, Experiment 3a and Experiment 3b.

Supplemental Table 1

| | **Finally determined CFS contrast** | **Finally determined target contrast** | **Breakthrough ratio** | |
|---|---|---|---|---|
| | | | **rewarded eye** | **non-rewarded eye** |
| S1 | 1 | 0.07 | 0.425 | 0.3312 |
| S2 | 1 | 0.09 | 0.3563 | 0.4562 |
| S3 | 1 | 0.15 | 0.2125 | 0.2125 |
| S4 | 1 | 0.2 | 0.3438 | 0.4625 |
| S5 | 1 | 0.04 | 0.5625 | 0.6188 |
| S6 | 1 | 0.18 | 0.6563 | 0.5 |
| S7 | 1 | 0.27 | 0.4562 | 0.475 |
| S8 | 1 | 0.8 | 0.4125 | 0.5313 |
| S9 | 1 | 0.45 | 0.2687 | 0.4 |
| S10 | 1 | 0.05 | 0.3812 | 0.3875 |
| S11 | 1 | 0.3 | 0.35 | 0.4313 |
| S12 | 1 | 0.1 | 0.4813 | 0.3812 |
| S13 | 1 | 0.4 | 0.575 | 0.475 |
| S14 | 1 | 0.4 | 0.425 | 0.4625 |
| S15 | 1 | 0.15 | 0.4188 | 0.3937 |
| S16 | 1 | 0.035 | 0.4688 | 0.5062 |
| S17 | 1 | 1 | 0.4063 | 0.425 |
| S18 | 1 | 0.1 | 0.45 | 0.4625 |

| | | | | |
|---|---|---|---|---|
| S19 | 1 | 0.1 | 0.4063 | 0.25 |
| S20 | 1 | 0.04 | 0.2375 | 0.275 |
| S21 | 1 | 0.04 | 0.5625 | 0.3688 |
| S22 | 1 | 0.1 | 0.5313 | 0.6062 |
| S23 | 1 | 0.15 | 0.5313 | 0.3438 |
| S24 | 1 | 1 | 0.5188 | 0.575 |
| S25 | 0.5 | 0.5 | 0.6563 | 0.4813 |
| S26 | 1 | 0.7 | 0.5313 | 0.4562 |
| S27 | 0.75 | 0.8 | 0.3812 | 0.5563 |
| S28 | 1 | 1 | 0.4437 | 0.4938 |
| S29 | 1 | 0.0375 | 0.5125 | 0.6125 |
| S30 | 1 | 0.2 | 0.2562 | 0.2938 |
| S31 | 1 | 0.1375 | 0.425 | 0.4375 |
| S32 | 1 | 0.4 | 0.5125 | 0.6062 |
| S33 | 1 | 0.02 | 0.2188 | 0.4063 |
| S34 | 1 | 0.3 | 0.275 | 0.1812 |
| S35 | 1 | 0.25 | 0.6 | 0.5437 |
| S36 | 1 | 1 | 0.2625 | 0.2313 |

Supplemental Table 2

| | Finally determined CFS contrast | Finally determined target contrast | Breakthrough ratio | |
|---|---|---|---|---|
| | | | rewarded eye | non-rewarded eye |
| S1 | 0.5 | 0.25 | 0.6062 | 0.55 |
| S2 | 0.5 | 0.9 | 0.5687 | 0.6438 |
| S3 | 1 | 0.4 | 0.5 | 0.5437 |
| S4 | 0.3 | 0.7 | 0.4625 | 0.4813 |
| S5 | 1 | 0.275 | 0.6813 | 0.4875 |
| S6 | 1 | 0.25 | 0.2562 | 0.4938 |
| S7 | 0.5 | 0.75 | 0.4437 | 0.45 |
| S8 | 1 | 0.35 | 0.2938 | 0.4437 |

| | | | | |
|---|---|---|---|---|
| S9 | 1 | 0.35 | 0.475 | 0.3563 |
| S10 | 0.45 | 1 | 0.5 | 0.6813 |
| S11 | 1 | 0.3 | 0.3187 | 0.2687 |
| S12 | 0.5 | 0.55 | 0.5062 | 0.4688 |
| S13 | 1 | 0.35 | 0.5 | 0.5188 |
| S14 | 0.7 | 1 | 0.475 | 0.475 |

Supplemental Table 3:

| | Finally determined CFS contrast | Finally determined target contrast | Breakthrough ratio of with-CFS trial | |
|---|---|---|---|---|
| | | | rewarded eye | non-rewarded eye |
| S1 | 1 | 0.9 | 0.4125 | 0.2875 |
| S2 | 1 | 0.4 | 0.2250 | 0.4625 |
| S3 | 1 | 0.3 | 0.4125 | 0.6125 |
| S4 | 1 | 0.8 | 0.2250 | 0.2750 |
| S5 | 1 | 0.16 | 0.3375 | 0.3875 |
| S6 | 1 | 0.15 | 0.1500 | 0.2375 |
| S7 | 1 | 0.04 | 0.5250 | 0.6000 |
| S8 | 1 | 0.15 | 0.3375 | 0.3000 |
| S9 | 1 | 0.65 | 0.3375 | 0.3375 |
| S10 | 1 | 0.7 | 0.4750 | 0.3375 |
| S11 | 1 | 0.7 | 0.4250 | 0.5375 |
| S12 | 1 | 0.15 | 0.5750 | 0.5000 |
| S13 | 1 | 0.1 | 0.1625 | 0.1500 |
| S14 | 1 | 0.6 | 0.4000 | 0.2500 |
| S15 | 1 | 0.6 | 0.4125 | 0.6000 |

Supplemental Table 4:

| | Finally determined CFS contrast | Finally determined target contrast | Breakthrough ratio | | | |
|---|---|---|---|---|---|---|
| | | | RO-RE | RO-NE | NO-RE | NO-NE |

| | | | | | | |
|---|---|---|---|---|---|---|
| S1 | 1 | 0.3 | 0.2125 | 0.175 | 0.525 | 0.475 |
| S2 | 1 | 0.4 | 0.5125 | 0.225 | 0.4 | 0.1375 |
| S3 | 1 | 0.25 | 0.2375 | 0.125 | 0.175 | 0.2 |
| S4 | 1 | 0.6 | 0.125 | 0.1875 | 0.2125 | 0.325 |
| S5 | 1 | 0.035 | 0.2667 | 0.5167 | 0.3833 | 0.4833 |
| S6 | 1 | 0.09 | 0.1667 | 0.3333 | 0.2167 | 0.2167 |
| S7 | 1 | 0.32 | 0.4625 | 0.225 | 0.225 | 0.25 |
| S8 | 1 | 0.16 | 0.475 | 0.25 | 0.3125 | 0.2125 |
| S9 | 1 | 0.25 | 0.3 | 0.4875 | 0.3375 | 0.45 |
| S10 | 1 | 0.18 | 0.2 | 0.3 | 0.2375 | 0.3125 |
| S11 | 1 | 0.55 | 0.5375 | 0.2125 | 0.325 | 0.125 |
| S12 | 1 | 1 | 0.4125 | 0.025 | 0.3625 | 0.05 |
| S13 | 1 | 0.07 | 0.1125 | 0.0125 | 0.2875 | 0.3625 |
| S14 | 1 | 0.15 | 0.1875 | 0.2125 | 0.2625 | 0.3 |
| S15 | 1 | 0.1 | 0.425 | 0.5125 | 0.35 | 0.4875 |
| S16 | 1 | 0.1 | 0.2167 | 0.25 | 0.4 | 0.4667 |
| S17 | 1 | 0.03 | 0.2833 | 0.2333 | 0.2833 | 0.5 |
| S18 | 1 | 0.1 | 0.275 | 0.35 | 0.525 | 0.4 |

Supplemental Table 5:

| | Finally determined CFS contrast | Finally determined target contrast | Breakthrough ratio | | | |
|---|---|---|---|---|---|---|
| | | | **RO-RE** | **RO-NE** | **NO-RE** | **NO-NE** |
| S1 | 0.65 | 0.6 | 0.1 | 0.075 | 0.3625 | 0.325 |
| S2 | 0.5 | 0.75 | 0.55 | 0.5875 | 0.575 | 0.7125 |
| S3 | 1 | 0.4 | 0.2875 | 0.425 | 0.325 | 0.4375 |
| S4 | 1 | 0.04 | 0.3 | 0.2375 | 0.2125 | 0.2 |
| S5 | 0.25 | 1 | 0.3125 | 0.2 | 0.125 | 0.6625 |
| S6 | 0.8 | 1 | 0.525 | 0.45 | 0.25 | 0.1 |
| S7 | 0.6 | 1 | 0.1875 | 0.275 | 0.3375 | 0.5125 |

| | | | | | | |
|---|---|---|---|---|---|---|
| S8 | 0.55 | 1 | 0.325 | 0.675 | 0.625 | 0.575 |
| S9 | 1 | 0.95 | 0.5375 | 0.7 | 0.5625 | 0.4625 |
| S10 | 0.18 | 1 | 0.375 | 0.3875 | 0.4125 | 0.5625 |
| S11 | 0.65 | 1 | 0.2875 | 0.3625 | 0.175 | 0.35 |
| S12 | 1 | 0.09 | 0.3875 | 0.5 | 0.4625 | 0.675 |
| S13 | 1 | 1 | 0.1625 | 0.475 | 0.25 | 0.3125 |
| S14 | 0.7 | 1 | 0.4375 | 0 | 0.55 | 0.05 |
| S15 | 0.5 | 1 | 0.725 | 0.625 | 0.5625 | 0.375 |
| S16 | 0.65 | 1 | 0.375 | 0.3875 | 0.375 | 0.5375 |
